# Cerebellar *Ex Vivo* Magnetic Resonance Imaging at its Feasibility Limit: Up to 77-Microns Isotropic Resolution using Low-Bandwidth Balanced Steady State Free Precession (LoBa-bSSFP) Sequences and 3T Standard Equipment

**DOI:** 10.1101/2024.04.18.589707

**Authors:** Matthias Weigel, Peter Dechent, Riccardo Galbusera, Erik Bahn, Po-Jui Lu, Ludwig Kappos, Wolfgang Brück, Christine Stadelmann, Cristina Granziera

## Abstract

**Background:** Ultra-high-resolution magnetic resonance imaging of the ex vivo brain is increasingly becoming an indispensable tool for studying the morphology and potential pathology of the brain. Despite the important role of the cerebellum in nervous system functions and motor control, as well as its potential damage in neurological diseases, it remains relatively understudied compared to other brain regions. One major reason is the even finer structures.

**Methods:** A balanced steady state free precession approach with receiver bandwidths as low as 50Hz/pixel and long repetition times of 36ms is suggested and optimized, called “LoBa-bSSFP”, which enhances the signal-to-noise ratio and alleviates strain on the gradient system for the ultra-high spatial resolutions. A radiofrequency phase cycle scheme is used to reduce potential artifacts. Only 3T MRI standard equipment is utilized for acquisition and basic image reconstruction of the ex vivo brain immersed in perfluoropolyether.

**Results:** The presented LoBa-bSSFP approach provides images with very good soft tissue contrast and a detailed visualization of cerebellar morphology. It enables isotropic resolutions of 98-microns for the entire cerebellum, a further refinement allows even up to 77-microns isotropic on a purely clinical MR system. The acquisitions preserved the integrity of the ex vivo cerebellum, so maintaining its connection to the cerebrum and brainstem.

**Conclusions:** Our findings demonstrate the feasibility of employing 3T based LoBa-bSSFP for true ultra-high-resolution ex vivo imaging of the cerebellum, reaching resolutions up to 77-microns isotropic and the potential to reveal subtle microscopic abnormalities of the cerebellar cortex. LoBa-bSSFP may be superior to conventional FLASH sequences in terms of acquisition efficiency and - in some cases - even contrast.

## Introduction

Ultra-high-resolution imaging (URI) of the human *ex vivo* brain has attracted a lot of interest lately ^1–12^. So far, less attention has been paid to the cerebellum, despite its importance for several nervous system functions in addition to fine-tuned motor control. Diseases such as Multiple Sclerosis (MS) have been shown to cause cerebellar damage. *Ex vivo* URI represents an excellent means for studying morphology and pathology of the human cerebellum.

Mainly due to reasons of SNR and therefore acquisition time challenges, *ex vivo* URI of the entire brain is mostly performed at 7T ^3–5,8^, although 3T MRI is more robust in terms of signal generation and 3T systems are more economical and widely available ^6,7,13–15^. Although earlier work on *ex vivo* MRI already studied balanced steady state free precession (bSSFP) sequences ^1,2,13^, in practice, RF-spoiled gradient echo sequences are typically the method of choice nowadays ^5,7,8,14^. Last year, we presented the idea of a low-bandwidth bSSFP (LoBa-bSSFP) approach for *ex vivo* MRI, using very low receiver bandwidths and long TRs for maximizing the SNR per unit time and reducing the strain on the MR gradient system ^15^. An RF phase cycle scheme was used to mitigate potential artifacts ^13,15^. As a result, a 115μm isotropic resolution was realized ^15^. Furthermore, different k-space sampling optimizations for accelerating imaging were tested on preliminary measurements ^15^.

In the following, we use an advancement of this LoBa-bSSFP approach to demonstrate the feasibility of high-quality 98μm and even 77μm isotropic acquisitions of the *ex vivo* cerebellum with standard 3T equipment. By taking advantage of a simple reduced FOV approach for the *ex vivo* cerebellum that we suggested in 2022, the integrity of the entire *ex vivo* brain is preserved, i.e., the cerebellum is still attached to the cerebrum and brainstem ^14^. So, the chosen concept is contrary to the imaging of tissue-blocks or slices ^16–19^, which can be performed in small-bore scanners using specialized receiver-coils. To reach the described feasibility limits, even more with standard 3T equipment, all URI acquisitions took place in the holiday and vacation time between Christmas Eve 2022 and New Year 2023.

## Materials and Methods

### Specimen preparation and experimental setup

The following experiments were approved by the ethical review committee of the Medical Center of Göttingen, Germany (29/9/10). All methods were carried out in accordance with the relevant guidelines and regulations. In further detail, clinical data and autopsy brain tissue were collected following established standard operating procedures (SOPs) developed within the MS Brain Bank of the KKNMS (German competence network for multiple sclerosis). The participant provided written informed consent for brain autopsy and for donation of his brain for research.

The *whole brain* of a deceased patient diagnosed with secondary progressive MS (male, 77 years old, disease duration 57 years) was fixed directly in 4% neutral buffered formaldehyde solution (formalin) approximately 24h after death.

For MRI acquisition, the brain was positioned in a 3D printed dome-shaped container ^20–22^ and immersed in the fluoropolymer perfluoropolyether (“Fomblin®”, Solvay Specialty Polymers USA, LLC, West Deptford, NJ, USA). A special circular surface plate kept the brain stable and avoided motion within the container. Residual air in the container was removed via degassing ports using a vacuum pump ^20–22^.

All investigations were performed on a clinical 3T whole-body MR system (Magnetom Prisma^Fit^, Siemens Healthineers, Erlangen, Germany). Only standard hardware components like the manufacturer-supplied 20-channel phased-array head and neck coil were employed. The URI experiments were conducted between Christmas Eve 2022 and beginning of January 2023, therefore harnessing the holidays and vacation time of Christmas and New Year 2022/2023 in Göttingen, Germany.

### LoBa acquisition approach

Our work is based on an advancement of the LoBa-bSSFP approach that we suggested last year ^15^, with preparatory work in ^13^. A reduced FOV focused on the cerebellum was realized via a simple but efficient approach we suggested in 2022 ^14^. As a result, the full reconstruction capability of the MR system allowed an 98μm isotropic acquisition of the entire cerebellum with LoBa-bSSFP. Based on a ‘clever positioning’ of the brain (cerebrum + cerebellum) and its container within the RF head and neck coil, a further reduction of the number of active coil elements freed additional reconstruction memory on the MR system, thus, allowing an increase to even 77μm isotropic.

Two dedicated cerebellum protocols, each operating at the described resolution limit, were set up. A LoBa-bSSFP approach with additionally programmed elliptical k-space sampling ensured the feasibility and stability of the measurements over hours and days: (1) 98μm isotropic: matrix 976×1080×576, TR/TE = 36.0ms/18.0ms, bandwidth = 50Hz/px, flip angle 40deg, eight different RF phase-cycles, 100 preparation pulses, acquisition time for one repetition TA1 = 6:21:05h, 21 repetitions, total acquisition time TAtotal ≈ 5.5 days. (2) 77μm isotropic: matrix 1248×1380×768, TR/TE = 36.0ms/18.0ms, bandwidth = 50Hz/px, flip angle 40deg, eight different RF phase-cycles, 100 preparation pulses, TA1 = 9:44:15h, 25 repetitions, TAtotal ≈ 10 days.

### Reconstruction of image data

For each phase-cycled repeated acquisition, both magnitude and phase images were reconstructed on-the-fly directly on the MR system with the standard postprocessing pipeline, consisting of the basic Fast Fourier Transform (FFT) and the standard intensity normalization filter (vendor name “prescan normalize”). Neither other kinds of filtering nor interpolation algorithms were utilized at this step. The images were exported into the standard DICOM format.

In an offline postprocessing step the exported DICOM images were converted into the NIfTI format using the free software *dcm2nix*. A rigid-body co-registration step with the free software *Elastix 14* followed ^23^. Afterwards the resulting images were complex-averaged.

Due to the immense size of 10GB per magnitude and phase volume scan (!) for the 77μm acquisitions, these measurements were split up into two half-volume parts, independently processed and combined again prior to co-registration.

The presented full-size and zoomed MR images in all figures were generated with the free software *ITK-SNAP 3*.*6*.*0* ^24^.

### Elective image postprocessing

A 3D Median filter and an adaptive 3D Median filter were tested as simple methods for image noise reduction.

## Results

Figures 1 to 4 illustrate that the developed LoBa-bSSFP approach is sustainable over the prolonged acquisition times of days needed for *ex vivo* URI of such – for MRI with standard clinical systems – extreme spatial resolutions up to 77μm isotropic. All acquisitions demonstrate a pronounced soft tissue contrast, the tightly folded thin layers of the cerebellar cortex and white matter interior were captured successfully. Noteworthy cerebellar structures such as the dentate nucleus are depicted well with their fine architecture (Figs. 1 and 3).

**Figure 1:**
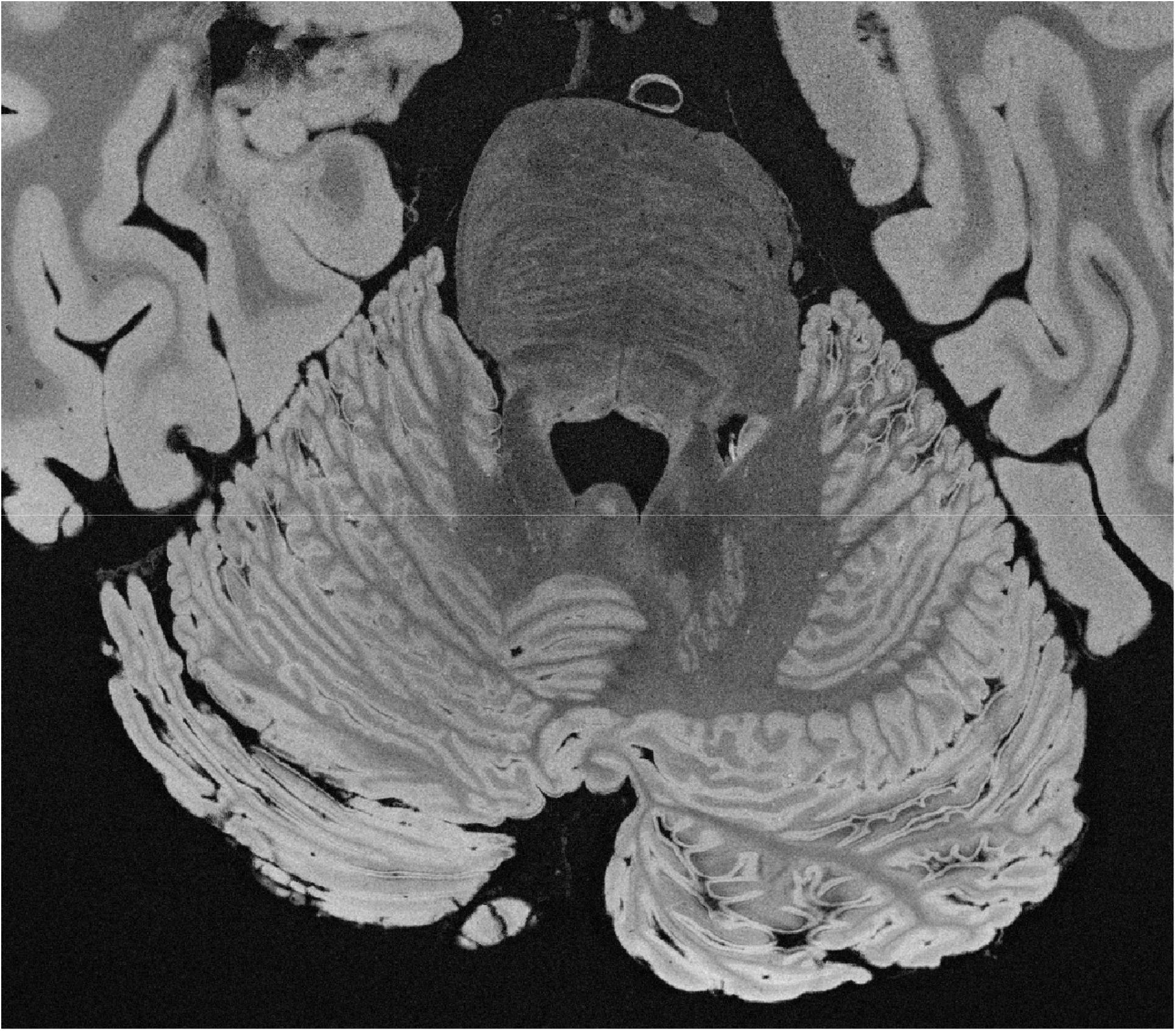
Representative slice of the cerebellum-focused 98μm full isotropic MRI acquisition, using the LoBa-bSSFP approach. The acquired entire FOV in the original transverse slice orientation is shown. The image unveils a rather impressing image quality of the intricate cerebellar fine structure.

**Figure 2:**
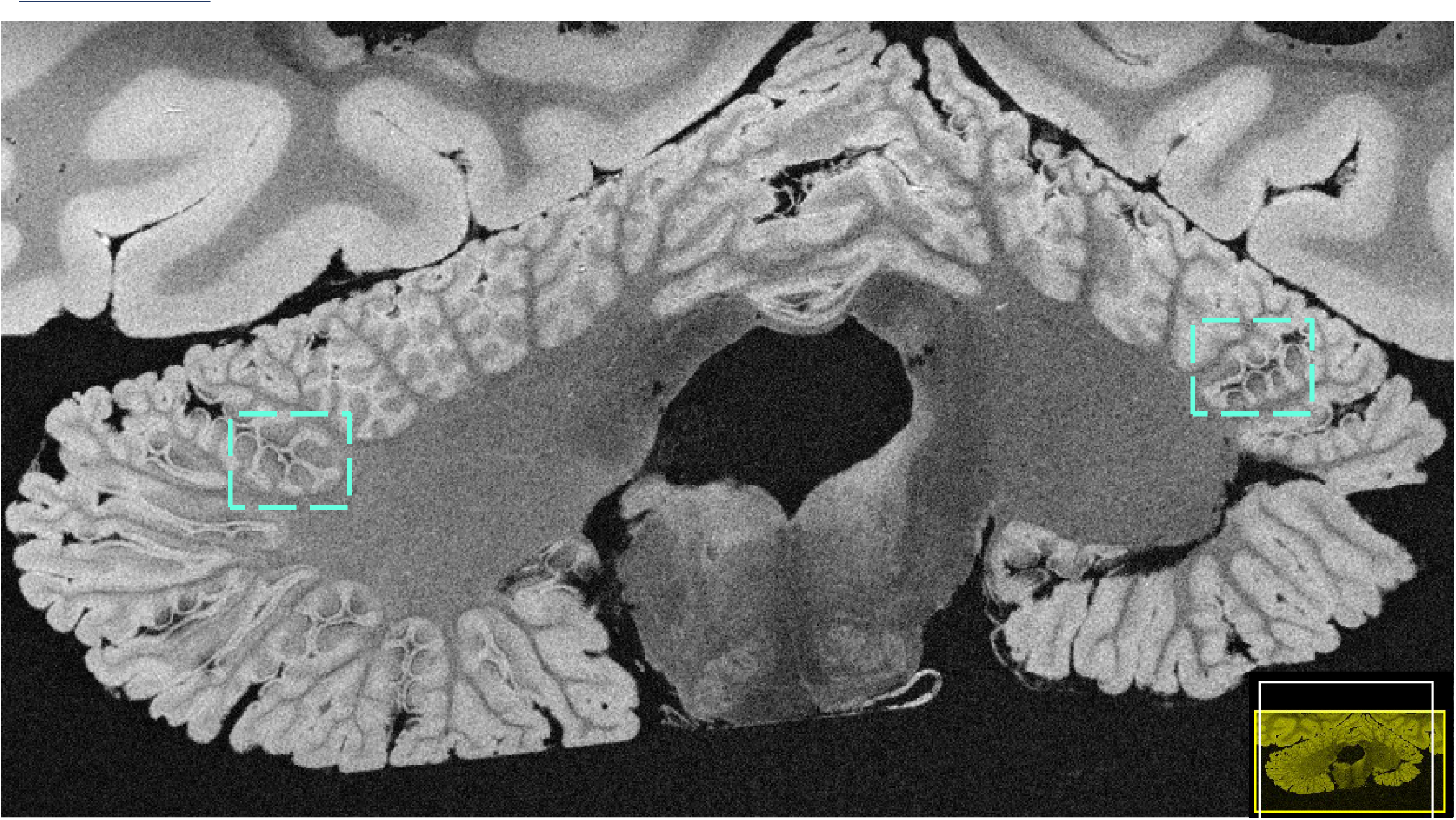
Coronal reformation of the 98μm LoBa-bSSFP measurement, underlining the full isotropic acquisition. The cerebellar foliae are shown in great detail, revealing focal symmetric microscopic alterations in the structure of the cerebellar cortex (see squares, for instance).

**Figure 3:**
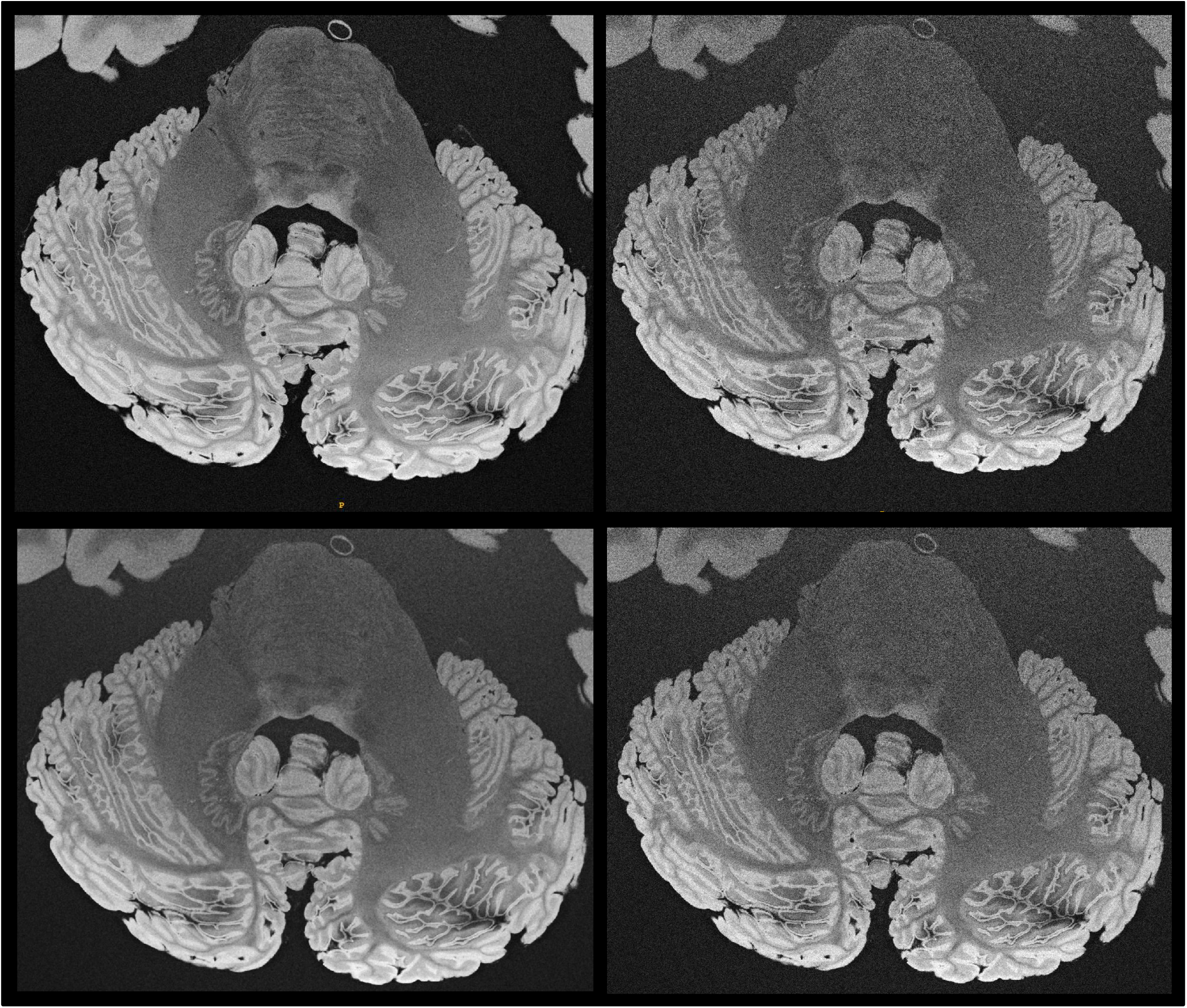
Top: Juxtaposition of the 98μm (left) with the 77μm acquisition (right). Whereas the depiction of the cerebellum seems to benefit from the increased resolution, the still reduced SNR seems to be counterproductive for resolving the fine structure of the brain stem. Bottom: Preliminary tests for noise reduction in the 77μm acquisition, using a median filter (left) and an adaptive median filter (right), the latter being a good compromise. Identical window-leveling was used for all images.

**Figure 4:**
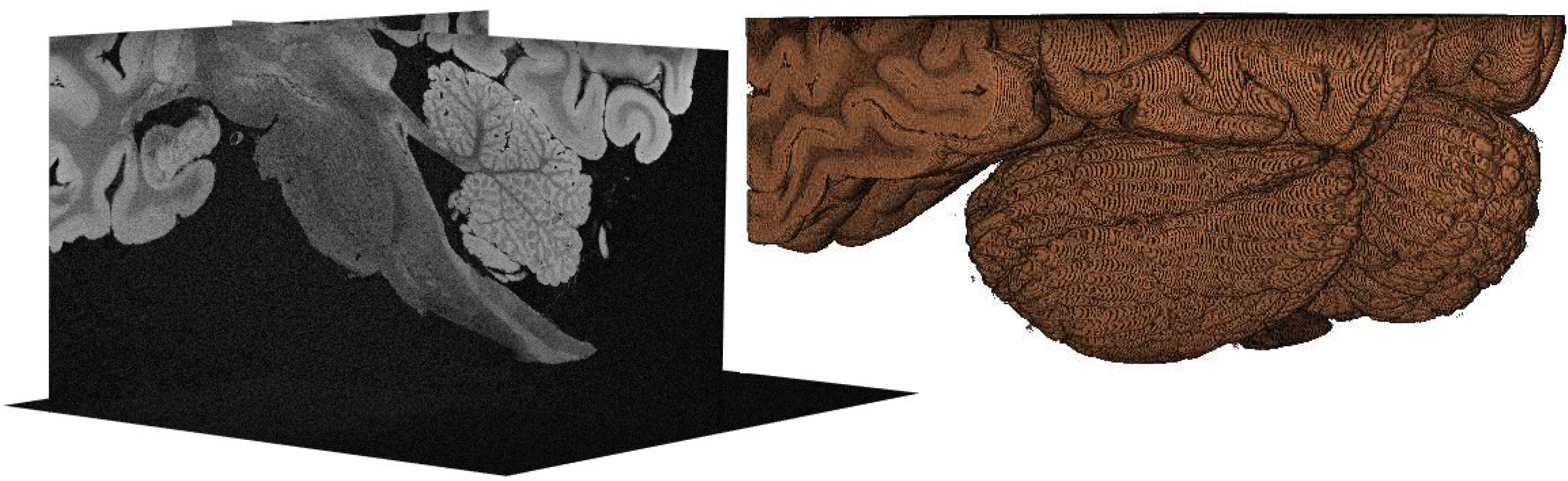
Slice planes view in 3D of the 98μm acquisition (left), depicting a sagittal view of both brainstem and cerebellum. The illustration represents a good means for visually estimating the acquired FOV. On the right side, a simple representative volume rendering was performed.

Particularly Figure 2, indicating several areas of focal alterations of the cerebellar cortex, underlines that LoBa-bSSFP image datasets provide the potential for resolving intricate pathological alterations associated with diseases such as MS.

## Discussion

The present work investigated the ‘ultimate boundaries’ for *ex vivo* URI of the cerebellum using a standard 3T MRI clinical system with from-the-shelf hardware components. Generally, the employed isotropic 3D acquisition approach is based on the idea of LoBa-bSSFP ^13,15^, which applies very low receiver bandwidths and long TRs for maximizing the SNR per unit time as well as reducing the permanent strain on the MR gradient system. Potential banding artifacts are effectively removed using an RF phase cycling scheme ^13,15^.

Recent publications emphasized the growing importance of MRI based *ex vivo* examinations for clinical neuroscience. Key factors for a successful depiction of morphology are ultra-high resolutions combined with homogeneous image intensities, a very good soft-tissue and pathology contrast as well as the lack of artifacts. As can be observed by Figs. 1 to 4, LoBa-bSSFP at 3T does fulfill these considerable constraints for cerebellar imaging – a rather impressive image quality is achieved. Utilizing a receiver bandwidth as low as 50 Hz/Px and a therefore necessary TR of 36ms, the chosen eight different RF phase cycles were sufficient to remove artifacts attributed to so called ‘banding’.

Total acquisition times of approximately 5.5 days and 10 days for the 98μm and 77μm isotropic resolution, respectively, may seem ‘outrageous’ and surely not be practical for most investigators. Although this criticism is probably true, it should be noted that the clear intent of this work was to test for the viable boundaries of cerebellar URI performed with 3T standard equipment and direct on-the-fly image reconstruction only. The purpose was to show how far one can get to obtain almost unprecedented MRI results. It should also be considered that spatial resolutions such as 77μm isotropic are usually found in the animal MRI system regime exclusively, at ultra-high fields but still with a much smaller total measurement volume.

The extensive acquisition times result from both the need for enhancing the SNR via signal averaging and from the necessity to spatially encode the desired volume in 3D with matrix sizes of roughly 1000^3^. In terms of SNR, a more dedicated RF coil with improved SNR could easily reduce the total acquisition time by a factor of 5, for instance. Moreover, in earlier work, we have shown that already a 115μm isotropic LoBa-bSSFP acquisition approach can provide exceptional results ^15^, however, it does have a considerably shorter acquisition time. As Fig. 3 also points out, already simple noise reduction methods like an (adapted) median filter can further improve image quality and may save acquisition time.

In one or the other way, all mentioned key-factors from above for high-quality images rely on a very good long-term stability of the acquisition approach. Compared to our extremely robust 3T URI-FLASH approach ^7,14^, a bSSFP based URI approach like LoBa-bSSFP is more drift-sensitive, however, this challenge could be tackled well with a dedicated image co-registration: Despite the lack of typical imaging artifacts, no blurring or smearing out of signal or tissue boundaries is observed, the edges are sharp. Naturally, the further increase in spatial resolution compared to ^7,14^ does also make the acquisition approach more drift-sensitive.

Although the results with LoBa-bSSFP based *ex vivo* acquisitions from here and ^15^ are more than rewarding, further work is needed in the link-up with histology, similar to Fig. 10 presented in ^7^. The cerebellum of the brain we analyzed in this work revealed the presence of numerous focal alterations of the cerebellar cortex (see Figs. 1 to 3).

To conclude, the presented long-term LoBa-bSSFP approach enables true *ex vivo* URI up to 77μm at 3T, thus providing a comprehensive microscopic view of the entire human cerebellum with the potential to reveal subtle microscopic abnormalities of the cerebellar cortex in MS and other neurological diseases.

## Relevance of the work

The presented LoBa-bSSFP approach facilitates fascinating insights into cerebellar morphology and potentially MS pathology. Particularly in the MRI based *ex vivo* ultra-high-resolution domain, the developed 3T bSSFP approach may be superior to conventional FLASH sequences in terms of acquisition efficiency and - in some cases - even contrast.

## List of abbreviations (alphabetical)

bSSFP: balanced SSFP
FLASH: fast low-angle shot
GM: gray matter
LoBa: Low-Bandwidth
URI: ultra-high-resolution imaging
WM: white matter

## Acknowledgements

We thank Heidelinde Brodmerkel, René Müller, and Katja Schulz for their very valuable technical assistance.

This work was funded by the Swiss National Science Funds PP00P3_176984 and PP00P3_206151 as well as supported by the German Ministry of education (BMBF; KKNMS German competence network for multiple sclerosis). Christine Stadelmann was supported by the Deutsche Forschungsgemeinschaft (DFG) transregional collaborative research center (CRC) TRR 274/1 and TRR274/2, Project ID 408885537 B01 and B03 and STA 1389/5-1 as well as the DFG under Germany’s Excellence Strategy (EXC 2067/1-390729940).

